# Characterization of liver-pancreas crosstalk following β-cell loss reveals a role for the molybdenum cofactor in β-cell regeneration

**DOI:** 10.1101/2024.04.09.588677

**Authors:** Christos Karampelias, Bianca Băloiu, Birgit Rathkolb, Patricia da Silva-Buttkus, Etty Bachar-Wikström, Susan Marschall, Helmut Fuchs, Valerie Gailus- Durner, Lianhe Chu, Martin Hrabě de Angelis, Olov Andersson

**Affiliations:** Department of Cell and Molecular Biology, Karolinska Institutet, Stockholm, Sweden; Institute of Diabetes and Regeneration Research, Helmholtz Munich, Neuherberg, Germany; Institute of Experimental Genetics, German Mouse Clinic, Helmholtz Zentrum München, Neuherberg, Germany; Institute of Molecular Animal Breeding and Biotechnology, Gene Center, Ludwig-Maximilians-Universität München, Munich, Germany; German Center for Diabetes Research (DZD), Neuherberg, Germany; Dermatology and Venereology Division, Department of Medicine, Karolinska Institutet, Stockholm, Sweden; Chair of Experimental Genetics, TUM School of Life Sciences, Technische Universität München, Freising, Germany; Department of Medical Cell Biology, Uppsala University, Sweden

## Abstract

Regeneration of insulin-producing β-cells is an alternative avenue to manage diabetes, and it is crucial to unravel this process in vivo during physiological responses to the lack of β-cells. Here, we aimed to characterize how hepatocytes can contribute to β-cell regeneration in a zebrafish model of β-cell ablation. Using lineage-tracing, we show that hepatocytes do not directly convert into β-cells even under extreme β-cell ablation conditions. A transcriptomics analysis of isolated hepatocytes following β-cell ablation displayed altered lipid- and glucose-related processes. Based on the transcriptomics, we performed a genetic screen that uncovers a potential role for the molybdenum cofactor (Moco) biosynthetic pathway in β-cell regeneration and glucose metabolism in zebrafish. Consistently, *Mocs2* haploinsufficiency in mice indicated dysregulated glucose metabolism and liver function. Together, our study sheds light on the liver-pancreas crosstalk and suggests that the molybdenum cofactor biosynthesis pathway should be further studied in relation to glucose metabolism and diabetes.

## Introduction

Diabetes is a multisystemic disease projected to affect up to 640 million people worldwide by 2040 (Sun et al., 2022). Insulin-producing pancreatic β-cells have a central role in diabetes pathophysiology. Patients with type 1 diabetes suffer from an autoimmune-mediated depletion of their β-cells while patients with type 2 diabetes manifest with reduced functional β-cell mass in later stages of the disease (Atkinson and Mirmira, 2023; Bakhti et al., 2019). Recent advances in diabetes research have shown intricate crosstalk between multiple tissues and β-cells that can regulate β-cell mass and function (Atkinson and Mirmira, 2023; Chiou et al., 2021; Kahraman et al., 2022; Shirakawa et al., 2017; Wang et al., 2019). Therefore, understanding how β-cell mass and function can be regulated by signals emanating from various tissues can increase our understanding of diabetes development, progression as well as generating novel therapies for the disease.

Stimulating β-cell regeneration holds interest as an approach to develop new therapeutics for diabetes patients. Important advances in understanding endogenous β-cell regeneration have been made possible using animal models. In these models, typically, there is an induced loss of β-cell mass that has led to studies into the mechanism of β-cell regeneration. A near-complete ablation of β-cells led to reprogramming of other pancreatic cell types including α-, δ-, γ- or acinar cells to β-cells (Carril Pardo et al., 2022; Chera et al., 2014; Druelle et al., 2017; Furuyama et al., 2019; Lee et al., 2018; Lu et al., 2016; Perez-Frances et al., 2021; Singh et al., 2022; Thorel et al., 2010; Ye et al., 2015). Proliferation of remaining β-cells also contributed to β-cell regeneration in less profound injury models (reviewed in (Basile et al., 2022). A controversial third regenerative pathway relies on pancreatic tissue-resident stem cell differentiation to endocrine fate, findings that are heavily debated for their applicability and can differ between injury models and model organisms (Delaspre et al., 2015; Ghaye et al., 2015; Gribben et al., 2021; Karampelias et al., 2022; Kopp et al., 2011; Magenheim et al., 2023; Sancho et al., 2014; Solar et al., 2009; Van de Casteele et al., 2013; Wang et al., 2020; Xiao et al., 2013; Xu et al., 2008; Zhao et al., 2021). Importantly, few of these studies have investigated how other organs and cell types outside the pancreas regulate regeneration of the β-cell mass.

Limited examples of signals from other tissues that contribute to β-cell regeneration exist (Shirakawa et al., 2017), and for example Adiponectin, an adipocyte-derived hormone, stimulated β-cell regeneration in mice (Ye et al., 2014). The liver is another central hub integrating molecular signals from pancreatic tissue (Titchenell et al., 2017). The crosstalk is evident as circulating factors from the liver can potentially regulate both α- and β-cell mass. First, a mouse model of hepatocyte-specific insulin receptor knockout showed an increase of β-cells (El Ouaamari et al., 2013; Michael et al., 2000). By utilizing this mouse model, Serpinb1 has been identified as one of the circulating factors that drives β-cell hyperplasia and correlating with insulin sensitivity (El Ouaamari et al., 2016; Glicksman et al., 2017). Second, additional in vivo models pointed to Igfbp1 as a secreted protein from the liver that promoted α- to β-cell transdifferentiation, kisspeptin as a liver-secreted neuropeptide that regulated insulin secretion from β-cells, and FGF21 as a liver-secreted cytokine that mediates β-cell regeneration in a mouse model treated with a glucagon receptor blocking antibody (Cui et al., 2023; Lu et al., 2016; Ouaamari et al., 2019; Song et al., 2014). Third, certain amino acids secreted from the liver promoted α-cell hyperplasia in mouse islets (Solloway et al., 2015). Fourth, *Pdx1* overexpression in mouse hepatocytes triggered a direct hepatocyte to β-cell conversion (Ferber et al., 2000; Meivar-Levy and Ferber, 2019). Taken together, these results suggest a promising but understudied role for the liver in stimulating β-cell regeneration under diabetic conditions.

In this study, we aimed to explore how the liver and more specifically the hepatocytes, contribute to β-cell regeneration in zebrafish. Zebrafish can regenerate most of its tissues following injury, and that is also true for their β-cell population (Goode et al., 2023). Here, we used the nitroreductase (NTR)-metronidazole (MTZ) zebrafish model to induce a near-complete β-cell ablation and study how the hepatocytes could be involved in the β-cell regeneration response (Curado et al., 2007; Pisharath et al., 2007). We showed that there is no spontaneous hepatocyte to β-cell conversion in the regenerating zebrafish larvae. Moreover, we characterized the transcriptome of zebrafish hepatocytes after β-cell ablation with the aim to identify immediate secreted signals or enzymes generating metabolites that can stimulate β-cell regeneration. We found that overexpression of the small isoform of the molybdenum cofactor synthesis 2 (Mocs2) enzyme (coding for the catalytic subunit of the molybdopterin synthase complex) in the hepatocytes could reduce glucose levels and stimulate a small but significant increase in β-cell regeneration. Mocs2 participates in the synthesis of the molybdenum cofactor (Moco) by catalyzing the synthesis of molybdopterin (Mayr et al., 2021), and we found that treatment with a molybdenum source, sodium molybdate, led to an increase in β-cell regeneration, albeit the phenotype was variable. In translating these findings to mice, we found that *Mocs2* haploinsufficient male mice manifested with a trend of impaired glucose tolerance. Together, our work has characterized important aspects of how the liver can affect β-cell regeneration and glucose homeostasis, and highlighted a role for Moco in this context.

## Results

### Characterization of the hepatocytes’ contribution to the spontaneous β-cell regeneration in zebrafish

The aim of this study is to characterize how the hepatocytes, which comprise the largest cell population of the liver, can contribute to pancreatic β-cell regeneration. We modelled β-cell loss using the NTR-MTZ ablation system in zebrafish. Briefly, in this transgenic zebrafish model the β-cells express the enzyme NTR. When the prodrug MTZ is added to the zebrafish water, NTR converts MTZ to a toxic byproduct and specifically ablates the β-cells. Using this β-cell ablation model, we explored the possibility that hepatocytes could be reprogrammed to β-cells, as documented previously, but without the forced expression of the *Pdx1* transcription factor (Meivar-Levy and Ferber, 2019). To this end, we used the *Tg(fabp10a:Cre);Tg(ubi:Switch)* zebrafish (Fig. 1A). In this lineage-tracing system, the *fabp10a*-expressing hepatocytes are permanently labelled by mCherry following recombination with the hepatocyte-specific Cre driver. We ablated the β-cells of zebrafish larvae with the addition of MTZ from 3 to 4 days post fertilization (dpf) and let the larvae regenerate for 2 days, a procedure which we from now on refer to as β-cell regeneration assay. We did not observe any hepatocyte-derived β-cells in the pancreas after 2 days of regeneration (Fig. 1B-B’). Moreover, there were no insulin-producing cells present in the liver with or without β-cell ablation (Fig. 1C-D’). These results demonstrate that hepatocytes are not spontaneously reprogrammed to β-cells in the highly regenerative developing zebrafish.

**Figure 1:**
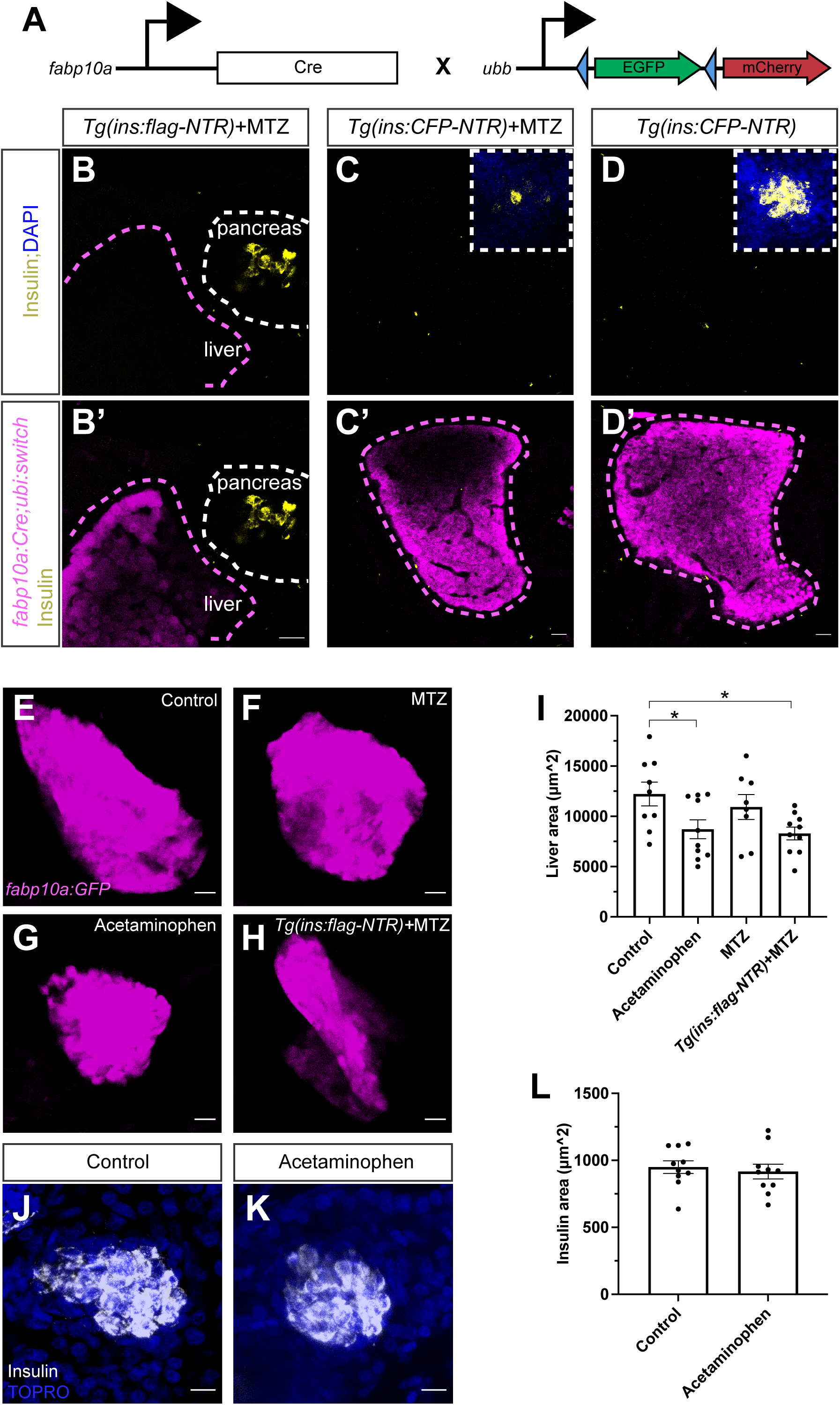
Hepatocytes’ contribution to the spontaneous β-cell regeneration in zebrafish. **A**: Schema showing the lineage-tracing approach to characterize hepatocyte to β-cell reprogramming using the *Tg(fabp10a:Cre)*;*Tg(ubi:Switch)* zebrafish . Blue arrowheads indicate the loxP sites. **B**: Single-plane confocal images of *Tg(fabp10a:Cre)*;*Tg(ins:flag-NTR);Tg(ubi:switch)* pancreas (**B**) and liver (**B’**) of 6 dpf zebrafish larvae after 2 days of β-cell regeneration and immunostained against insulin. White dashed line outlines the pancreas and the magenta dashed line outlines the border of the liver. Scale bar, 20 µm. *n* ≥ 10 larvae examined from two independent experiments. **C**-**D**: Single-plane confocal images of liver and primary islets of *Tg(fabp10a:Cre)*;*Tg(ins:CFP-NTR);Tg(ubi:switch)* 6 dpf larvae, with (**C**-**C’**) or without (**D**-**D’**) MTZ treatment, i.e. β-cell ablation. Larvae in (**C**-**C’**) were left to regenerate their β-cells for 2 days. White dashed line outlines the insets of the primary islet of the pancreas (**C**,**D**) and the magenta dashed line outlines the border of the liver (**C’**,**D’**). Scale bar, 20 µm. *n* ≥ 10 larvae examined from two independent replicates. **E**-**I**: Maximum projections of livers from control (**E**), MTZ-treated (**F**), acetaminophen-treated (**G**) and *Tg(ins:flag-NTR)*+MTZ-treated (**H**) *Tg(fabp10a:GFP)* 4 dpf zebrafish larvae. Chemical treatments were carried out at 3-4 dpf. Quantification showed a significant decrease of the hepatocyte area following hepatocyte damage or β-cell ablation (**I**). Scale bar, 20 µm. *n* = 8-10. Data are presented as mean values ± SEM. One-way ANOVA was used to estimate statistical significance followed by a Holm-Šidák multiple comparison test. \**P* = 0.0325 (control vs acetaminophen); \**P* = 0.0237 (control vs *Tg(ins:flag-NTR)*+MTZ). **J**-**L**: Representative maximum projections of pancreatic islets in control (**J**) and acetaminophen-treated (3-4 dpf) (**K**) 5 dpf zebrafish larvae immunostained against insulin. Nuclei were counterstained with TO-PRO-3. Quantification of the insulin area is shown in (**L**). Scale bar, 10 µm. *n* = 10.

Next, we aimed to damage the hepatocytes and assess if there is a reduction in the basal β-cell regeneration in the pancreas of zebrafish larvae. To this end, we treated the zebrafish larvae with a high concentration of acetaminophen, a drug that has previously been shown to induce hepatocyte damage in zebrafish (North et al., 2010). We confirmed that 24 hours of acetaminophen treatment reduced liver size in *Tg(fabp10a:GFP)* zebrafish larvae. We observed that β-cell ablation reduced liver size to comparable levels as acetaminophen treatment (Fig. 1E-I). This effect was independent of MTZ treatment as larvae without the *Tg(ins:flag-NTR)* transgene but treated with MTZ had normal liver size. Conversely, we explored whether hepatocyte damage/acetaminophen treatment impaired β-cell development; Acetaminophen treatment for 24 hours did not affect β-cell development of 5 dpf zebrafish (Fig. 1J-L). We corroborated these results with a genetic model of hepatocyte ablation based on the same NTR-MTZ model. Transgenic *Tg(fabp10a:CFP-NTR)* zebrafish larvae were treated with MTZ to ablate almost all hepatocytes, as shown before (Choi et al., 2014). Similar to acetaminophen treatment, this did not lead to any changes in β-cell development (Supplementary Fig. 1). Overall, our data shows that no hepatocyte to β-cell reprogramming is observed in zebrafish larvae and that liver damage does not affect β-cell development.

### Transcriptomic changes in hepatocytes following β-cell ablation

Secreted proteins from the liver have been shown to contribute to compensatory β-cell expansion in zebrafish and mice. Therefore, we decided to explore the possibility that additional hepatocyte-derived secreted signals can stimulate β-cell expansion. To this end, we characterized the transcriptome of 2-month-old zebrafish hepatocytes following β-cell ablation. In this experiment, we ablated β-cells in our zebrafish model followed by one day of regeneration, and then sacrificed the zebrafish and dissected out the livers. We used the *Tg(fabp10a:GFP)* as the control samples and *Tg(fabp10a:GFP);Tg(ins:flag-NTR)* zebrafish as the β-cell ablated samples. Then, we FACS the GFP^+^ hepatocytes, extracted RNA and performed bulk RNA-Seq (Fig. 2A). Further, we confirmed that the β-cell ablation was successful by measuring the blood glucose levels at the time of sacrificing the zebrafish (Fig. 2B).

**Figure 2:**
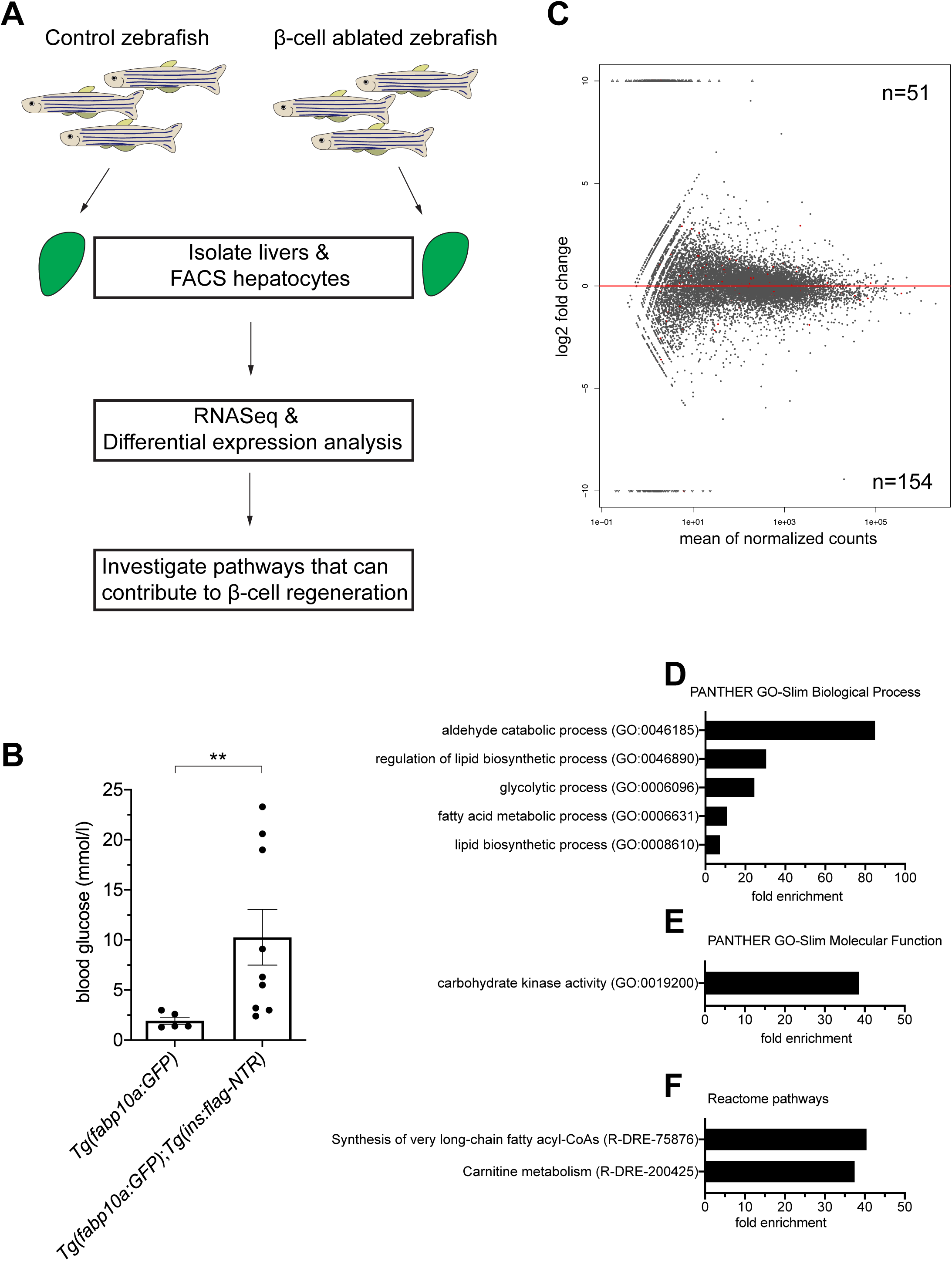
Transcriptomic changes in hepatocytes following β-cell ablation. **A**: Schema showing the experimental design to identify transcriptional changes in hepatocytes following β-cell ablation in 2-month old zebrafish. **B**: Glucose measurements of the 2-month old zebrafish used for hepatocyte isolation. Measurements were made before liver dissection. *n* = 5-9. Mann-Whitney test was used to assess significance. \*\**P* = 0.0045. **C**: Log2 fold change plot showing the upregulated and downregulated genes in hepatocytes following β-cell ablation. The significantly differentially expressed genes (padj < 0.05) are highlighted as red dots. **D**-**F**: Statistical overrepresentation analysis of the downregulated genes from (**C**) using the PantherDB tool. The fold enrichment of the significantly enriched processes (**D**), molecular function (**E**) and reactome pathways (**F**) are shown respectively.

We performed differential expression analysis to characterize the transcriptomic changes in the hepatocytes following β-cell ablation. Overall, we observed a greater number of downregulated genes compared to upregulated genes (Fig. 2C). Key genes that have previously been shown to be regulated by insulin were downregulated in hepatocytes following β-cell ablation (Supplementary file 1). For example, glucokinase (*gck*) was the most significantly downregulated gene in our dataset. Moreover, we observed a non-significant upregulation of both *serpinb1* and *igfbp1a*, two secreted peptides that were previously been implicated in β-cell regeneration (Supplementary Fig. 2A). Subsequently, we performed a gene ontology enrichment analysis to reveal the pathways that were significantly affected in hepatocytes following β-cell ablation. In general, cellular processes and metabolic processes were two of the most affected biological processes in both up- and downregulated genes (Supplementary Fig. 2B-C). We next performed a statistical overrepresentation analysis and found no significantly enriched gene-categories for the upregulated genes. On the contrary, glycolytic and lipid related biological processes were significantly enriched in the downregulated genes (Fig. 2D). In addition, downregulated genes were enriched in carbohydrate kinase activity as well as in the synthesis of very long-chain fatty acyl-CoAs and carnitine metabolism reactome pathways (Fig. 2E-F). Overall, our transcriptomic characterization of hepatocytes following β-cell ablation in 2-month-old zebrafish showed a downregulation of glycolytic and lipid-related metabolic processes.

### An in vivo genetic screen to identify hepatocyte genes that can enhance β-cell regeneration

Next, we focused our analysis on the significantly upregulated genes in the RNA-Seq dataset. We aimed to determine if the upregulated genes coding for secreted proteins or enzymes could accelerate β-cell regeneration when overexpressed in hepatocytes to supra-physiological levels. Our dataset contained two genes coding for secreted proteins and eleven genes coding for enzymes that were significantly upregulated (p adj<0.05) (Fig. 3A). We cloned these genes under the control of the liver-specific *fabp10a* promoter and injected them in the *Tg(ins:kaede);Tg(ins:CFP-NTR)* zebrafish model to perform our standard β-cell regeneration assay. Of note, we did not perform the regeneration assay with the *igfbp5a* gene as we tested it in our previously published work (Lu et al., 2016). Mosaic overexpression of the single upregulated gene coding for a secreted protein, *sdf2l1*, did not increase β-cell regeneration (Fig. 3B). Subsequently, we overexpressed the nine of the eleven significantly upregulated enzymes (as we did not manage to clone two of the enzymes, namely *pdzrn3a* and *tor1l1*, see methods) and discovered a modest, yet significant, increase in β-cell regeneration upon overexpression of the small isoform of the *mocs2* gene. On the contrary, overexpressing the big isoform of the *mocs2* enzyme did not result in a similar phenotype, suggesting different functions of the two protein isoforms (Fig. 3C). Overall, our genetic screen pointed to the small isoform of *Mocs2* as a gene candidate that could be implicated in β-cell regeneration via its expression in the liver.

**Figure 3:**
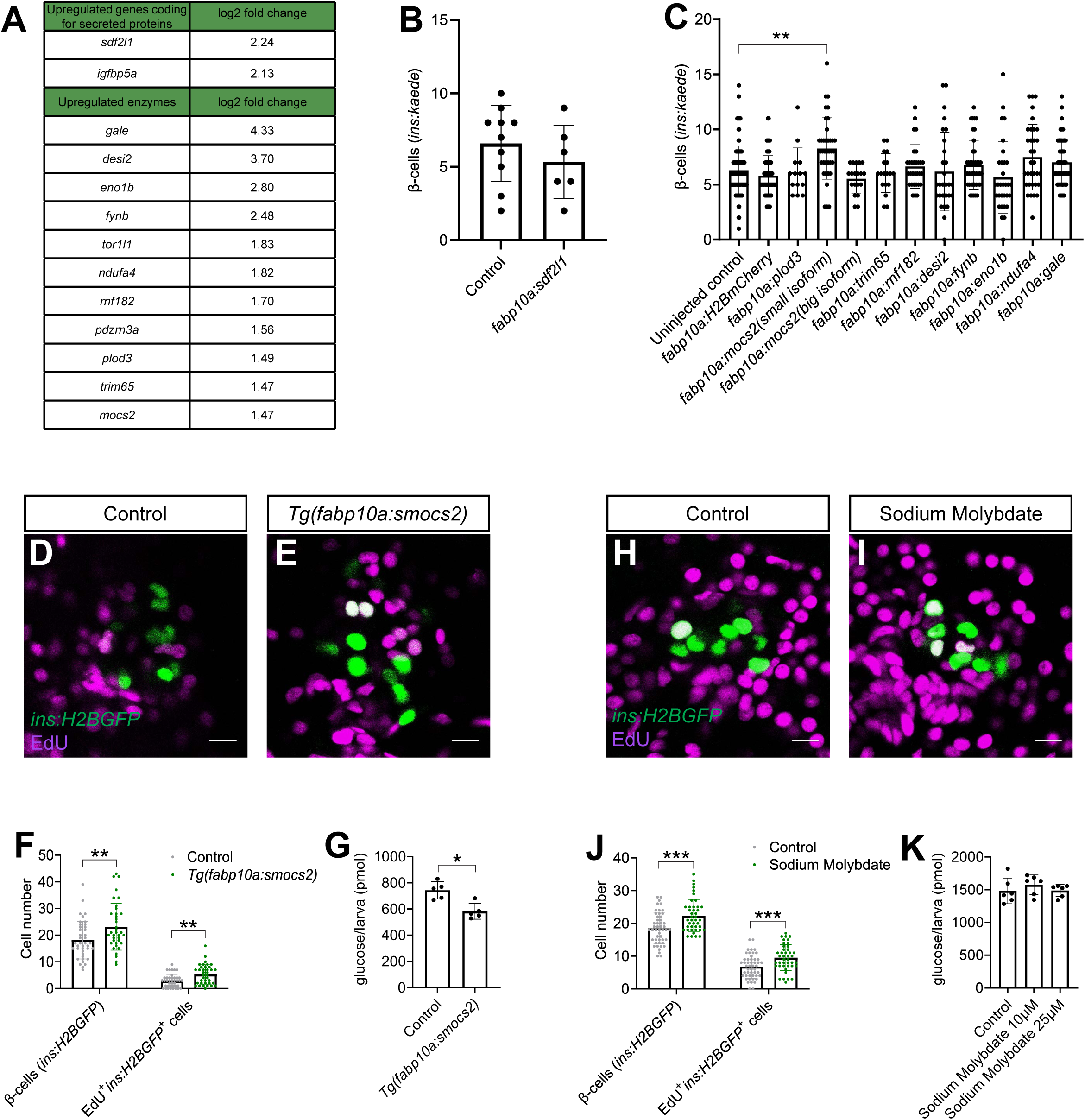
Genetic screen reveals a role for the molybdenum cofactor biosynthetic pathway in β-cell regeneration. **A**: Table showing the log2 fold changes of the significantly upregulated genes encoding for secreted proteins and enzymes in hepatocytes following β-cell ablation. **B**: Quantification of β-cells from 6 dpf *Tg(ins:kaede)*;*Tg(ins:CFP-NTR)* larvae overexpressing *sdf2l1* in hepatocytes. *n* = 6-10. Data are presented as mean values ± SEM. **C**: Quantification of β-cells in control and zebrafish larvae injected at the one-cell stage with vectors driving expression of the enzymes identified in hepatocytes (**A**) under the control of the *fabp10a* promoter (together with mRNA encoding the transposase enzyme to induce genomic integration). Following injections, β-cells were ablated from 3 to 4 dpf and the β-cells were counted manually after two days of regeneration in the *Tg(ins:kaede)*;*Tg(ins:CFP-NTR)* zebrafish larvae at 6 dpf. *n* = 14-104. Data for the control experiments were pooled from 4 independent experiments. If there was a positive hit in the first experiment, the experiments were repeated and data pooled in this graph. Kruskal-Wallis test followed by Dunn’s multiple comparison test was used to assess statistical significance. \*\**P* = 0.0038. Data are presented as mean values ± SEM. **D**-**F**: Single-plane confocal images of pancreatic islets of control (**D**) and *Tg(fabp10a:smocs2)* (**E**) *Tg(ins:flag-NTR)*;*Tg(ins:H2BGFP)* 6 dpf zebrafish larvae after two days of β-cell regeneration, during which EdU incubation occurred. Quantification of the number of *ins:H2BGFP^+^ and ins:H2BGFP^+^*EdU*^+^* cells (**F**). Scale bar, 10 µm. *n* = 37-40. Mann-Whitneytest was used to assess significance. \*\**P* = 0.0079 and 0.0037 respectively. Data are presented as mean values ± SEM and are pooled from 3 independent experiments. **G**: Glucose levels in *Tg(ins:flag-NTR);Tg(fabp10a:smocs2)* 6 dpf larvae. Four larvae were pooled for each replicate. *n* = 5. Mann-Whitney test was used to assess significance. \**P* = 0.0159. Data are presented as mean values ± SEM. **H**-**J**: Single-plane confocal images of pancreatic islets of control (**H**) and sodium molybdate treated (**I**) *Tg(ins:flag-NTR)*;*Tg(ins:H2BGFP)* 6 dpf zebrafish larvae after two days of β-cell regeneration, during which EdU incubation occurred. Quantification of the number of *ins:H2BGFP^+^ and ins:H2BGFP^+^*EdU*^+^* cells (**J**). Scale bar, 10 µm. *n* = 42-49. Mann-Whitney test was used to assess significance for β-cell numbers and unpaired Student’s t-test was used to assess significance for *ins:H2BGFP^+^*EdU*^+^*. \*\*\**P* = 0.0005 and 0.0008 respectively. Data are presented as mean values ± SEM and are pooled from 3 independent experiments. **K**: Glucose levels of *Tg(ins:CFP-NTR)* 6 dpf larvae treated with 10 µM or 25 µM sodium molybdate. Four larvae were pooled for each replicate. *n* = 6. Data are presented as mean values ± SEM.

### Phenotypic characterization of *Mocs2* overexpression in zebrafish

Following our genetic screen, we generated a stable transgenic line overexpressing the small isoform of *mocs2* in hepatocytes, named from here on *Tg(fabp10a:smocs2)*. We observed that the *Tg(fabp10a:smocs2)* larvae increased β-cell regeneration similar to the mosaic overexpression (Fig. 3D-F). The extent of β-cell regeneration was variable between experimental replicates with some showing a profound increase in β-cell regeneration while other experiments showed a less pronounced phenotype that did not reach statistical significance. To study the cellular mechanism behind the observed β-cell regeneration, we incubated control and *Tg(fabp10:smocs2)* larvae with EdU during the regenerative period to assess β-cell proliferation. We observed a cumulative significant increase of EdU incorporation in regenerating β-cells, yet the phenotype was variable between replicates similar to the β-cell regeneration observation (Fig. 3D-F). This suggests that there are other biological factors that need to coordinate with the *mocs2* overexpression to reach the full potential of the phenotype. Additionally, we assessed the glucose levels in the *Tg(fabp10a:smocs2)* larvae. We noticed a decrease of glucose levels in larvae on the *Tg(fabp10:smocs2)* background compared to control larvae following β-cell ablation, a phenotype that was more consistent than the increase in β-cell regeneration (Fig. 3G).

Next, we aimed to stabilize and clarify the effect of Moco biosynthesis in β-cell regeneration using a non-genetic approach. To this extent, we treated zebrafish larvae with sodium molybdate, which has previously been used as a source of molybdenum in *E. coli* cultures and can rescue defects in the Moco biosynthetic pathway (Warnhoff and Ruvkun, 2019), for two days following β-cell ablation. Sodium molybdate increased β-cell regeneration through increased β-cell proliferation similarly to the overexpression of the small isoform of *mocs2*, yet the phenotype also appeared to be variable resembling the genetic model (Fig. 3H-J). On the contrary, none of the experiments with sodium molybdate treatments showed decreased glucose levels highlighting a difference between the genetic and chemical approach (Fig. 3K).

Overall, our combinatorial chemical/genetic approach to perturb the Moco biosynthetic pathway showed a role of this pathway in β-cell biology and glucose homeostasis, but additional factors are needed to recapitulate a more robust phenotype.

### Gene expression of the core genes of Moco biosynthetic pathway across species

To better understand the Moco pathway, we assessed the expression levels of the genes involved in Moco biosynthesis as well as the enzymes that use molybdenum cofactor for their function across species. In zebrafish, we identified orthologues of all the mammalian genes involved in Moco biosynthesis, namely *mocs1*, *mocs2*, *mocs3*, *gphna*, *gphnb* together with homologues of enzymes requiring molybdenum for their function including *suox*, *xdh*, *aox5* and *aox6*. All enzymes involved in Moco biosynthesis and utilization were expressed in our RNASeq dataset of isolated hepatocytes except for *gphnb* (the paralogue of *ghpna*) and *aox5* (Supplementary Fig. 3A). Comparing how their expression was affected by β-cell ablation, we observed that only *mocs2* was significantly upregulated with the rest of the enzymes’ expression not significantly affected (Supplementary Fig. 3B). We then assessed the expression levels of the same enzymes in our RNASeq of isolated islets from zebrafish larvae with or without β-cell ablation, data that we previously published (Karampelias et al., 2021). We observed a similar expression level for all enzymes of the pathway except *gphnb,* which had expression below the level of detection, similar to its absence of expression in hepatocytes (Supplementary Fig. 3C). Further, β-cell ablation did not significantly upregulate the expression of any genes in the pathway, with the most noticeable change being a downregulation of the *suox* enzyme (Supplementary Fig. 3D). Expression levels of *ins* was used as a reference for both datasets (Supplementary Fig. 3A-D).

We used publicly available single-cell RNASeq (scRNA-Seq) datasets and atlases to assess the expression of the genes involved in the molybdenum cofactor biosynthetic pathway in mice and human tissues. Using a recently published mouse pancreas atlas (Hrovatin et al., 2023), we observed that *Mocs2* was the highest expressed gene of the pathway across the different cell types of the pancreas (Supplementary Fig. 4A). Expression levels did not vary in different models of diabetes in the same atlas (Supplementary Fig. 4B). The *Suox* enzyme had its highest expression in the endocrine part of the pancreas, albeit at low levels. Compared to the mouse pancreas atlas, we enquired a recent human scRNA-Seq compilation dataset including 65 pancreata from human donors with/without diabetes from the Human Pancreas Analysis Program (Elgamal et al., 2023). Similar to the mouse pancreas, *MOCS2* had the highest expression across most pancreatic cell types, with highest expression being in β-cells and cycling α-cells (Supplementary Fig. 5A-G). On the contrary, *MOCS2* had low basal expression in human liver cell types, while *GPHN*, *AOX1* and *XDH* had the highest expression in hepatocytes in a recently published human liver scRNA-Seq atlas (Supplementary Fig. 5H-N) (Wu et al., 2023). Overall, we observed a high gene expression of Mocs2 in mouse and human endocrine pancreas compared to the rest of the genes of the Moco biosynthetic pathway.

### Phenotypic characterization of *Mocs2* mutant mice

To examine if any of the molybdenum-related phenotypes observed in the zebrafish model could be translated to mice, we used the deep phenotyping of a *Mocs2* mutant mouse line generated as part of the international mouse phenotyping consortium (Groza et al., 2023). Homozygous Mocs2 mutants were embryonically lethal showcasing the importance of this pathway for proper embryonic development. Therefore, we assessed if any metabolic phenotypes were present in the heterozygous mice (*Mocs2^+/-^*) between 12 and 16 weeks of age. Pancreatic, intestinal, stomach (data not shown) and liver morphology, as evaluated in hematoxylin-eosin stained sections was unchanged between wild-type and *Mocs2^+/-^* (Supplementary Fig. 6). In the endocrine pancreas, immunohistochemical staining of insulin and glucagon demonstrated normal protein patterns and correct mouse islet morphology (Supplementary Fig. 6C&F). We further characterized the terminal differentiation of α-cells and β-cells using the Ucn3 immunostaining that marks mature mouse β-cells (Blum et al., 2012). Regardless of sex and genotype, Ucn3 was absent from α-cells suggesting correct lineage allocation (Fig. 4A-H). Overall, no obvious morphological abnormality was observed in the islets (Fig. 4A-H and Supplementary Fig. 6). Then, we assessed metabolic and glucose phenotypes of these mice in both sexes. There were no differences in the body weight between wild-type or *Mocs2^+/-^* mice (Fig. 4I-K). Overall, *Mocs2^+/-^* mice showed a trend towards higher fasting glucose as well as signs of glucose intolerance indicated by slightly delayed glucose clearance in an intraperitoneal glucose tolerance test (IPGTT) (particularly in male mice), but the results did not reach statistical significance (Fig. 4M,P). This is due to high variability in the glucose measurements of the *Mocs2^+/-^* similar to the observed phenotypes of the zebrafish larvae. No phenotypic changes were observed in female *Mocs2^+/-^* mice suggesting that there might be sex-specific glucose effects (Fig. 4L-Q). Fasting glucose levels and IPGTT approached statistical significance when the results from both sexes were combined (Fig. 4N-Q). Lastly, we also examined global markers of liver cell malfunction including alanine aminotransferase activity (ALAT), bilirubin and alkaline phosphatase activity (ALP). Similar to the glucose phenotype, male mice appeared to have altered biochemical profiles for total bilirubin levels and ALP activity (Supplementary Fig. 7). Increased ALP activity in males could signal liver or bone defects in the *Mocs2^+/-^* mice (Supplementary Fig. 7H), while reduced bilirubin levels are harder to interpret (Supplementary Fig. 7E). A summary of all the clinical parameters assessed for this experiment can be found in Supplementary File 2. Overall, the deep phenotyping of the *Mocs2^+/-^* mice showed a trend towards sex-specific (male mice) altered glucose metabolism and liver alterations, in agreement to the variable phenotype observed in the zebrafish model.

**Figure 4:**
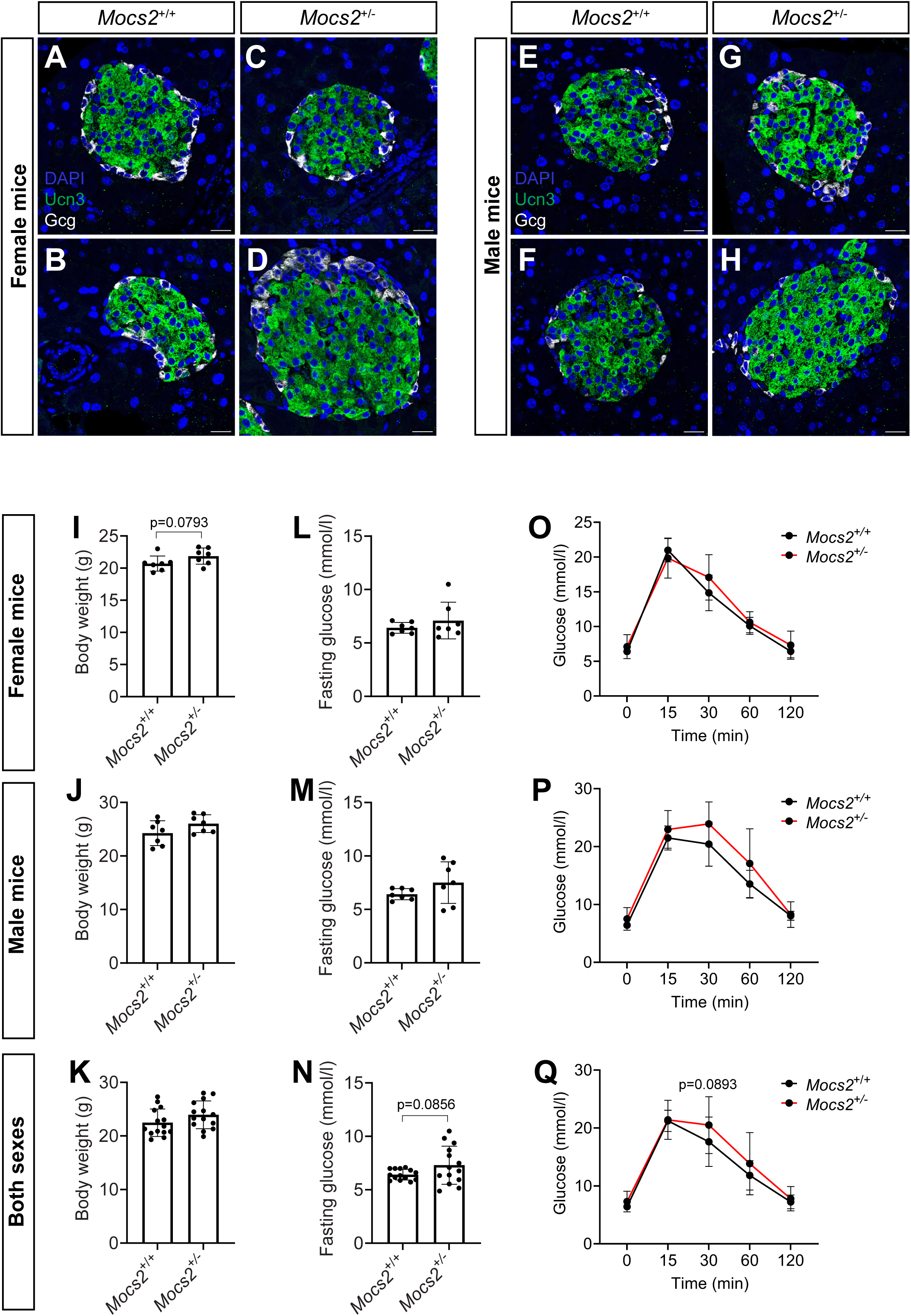
Phenotyping of *Mocs2^+/-^* mice. **A**-**H**: Single-plane confocal images of mouse islets from female *Mocs2^+/+^* (**A**-**B**) and *Mocs2^+/-^* (**C**-**D**), as well as male *Mocs2^+/+^* (**E**-**F**) and *Mocs2^+/-^* (**G**-**H**), stained for Gcg, Ucn3 and counterstained with DAPI. Scale bar 20 µm. **I**-**K**: Body weight measurement before the start of the fasting period for the IPGTT from female (**I**), male (**J**) and combined data (**K**) for *Mocs2^+/+^* and *Mocs2^+/-^* mice. Mann-Whitney test was used to assess significance. **L**-**N**: Fasting glucose measurement from female (**L**), male (**M**) and combined data (**N**) for *Mocs2^+/+^* and *Mocs2^+/-^* mice. Student’s t-test was used to assess significance. **O**-**Q**: IPGTT results from female (**O**), male (**P**) and combined data (**Q**) for *Mocs2^+/+^* and *Mocs2^+/-^* mice. Two-way ANOVA followed by Šidak’s multiple test was used to assess significance.

## Discussion

In this work, we aimed to characterize the liver-pancreas crosstalk and how this could affect β-cell regeneration in the highly regenerative zebrafish model. Our results indicate that there is no direct reprogramming of hepatocytes to β-cells in vivo. Previous reports that showed these reprogramming events were based on overexpression of the fate determinant transcription factor *Pdx1* in mice (Ber et al., 2003; Ferber et al., 2000). It was also reported that the ductal cells of the liver could be reprogrammed to β-cells upon genetic manipulation in mice (Banga et al., 2012). It needs to be noted that the ductal system of the liver and the pancreas derives from a common progenitor and that the ductal cells of the pancreas are known to contribute to β-cell regeneration in zebrafish (Delaspre et al., 2015; Delous et al., 2012; Ghaye et al., 2015; Karampelias et al., 2021; Karampelias et al., 2022; Liu et al., 2018; Manfroid et al., 2012; Mi et al., 2023). Whether or not the intrahepatic duct could differentiate into β-cells in zebrafish remains to be seen.

As part of this work, we characterized the transcriptome of isolated hepatocytes from 2-month-old zebrafish after β-cell ablation. Our data revealed that glucose and lipid metabolism are the most affected biological processes in the absence of β-cells/insulin and associated increased glucose levels. This data confirms the role of insulin as a master regulator of metabolism in the zebrafish liver and stresses the conservation of its action to the mammalian orthologue (Saltiel, 2021). However, the number of genes significantly changed was not dramatic with only 51 significantly upregulated genes discovered in our analysis, suggesting that perhaps more time is needed following β-cell ablation for additional genes to be upregulated. This could be attributed to low statistical power due to the small number of biological replicates of our study, which is a potential limitation of our work. This observation is strengthened by the fact that both *serpinb1* and *igfbp1a*, genes encoding proteins that were previously implicated in β-cell regeneration and secreted from the liver, were non-significantly upregulated in our dataset (El Ouaamari et al., 2016; Lu et al., 2016). Only two of the upregulated genes had the potential to code for circulating proteins but none of them stimulated β-cell regeneration when overexpressed.

We also performed a genetic screen by overexpressing the upregulated enzymes in the hepatocytes. We reasoned that in this way we could affect the level of a circulating metabolite that might stimulate β-cell regeneration and/or glucose levels. Our screen revealed a role for the small isoform of *mocs2* in β-cell regeneration and lowering of glucose. Mocs2 is an enzyme involved in the cascade responsible for the generation of the molybdopterin metabolite and intermediate of the molybdenum cofactor that donates the metal molybdenum to enzymes that need it for their action, including sulfite oxidases and xanthine oxidoreductases. The *mocs2* gene is bicistronic and two isoforms are transcribed. The big and the small isoforms come together to form a heterotetrameric complex responsible for the generation of molybdopterin (Mayr et al., 2021). We observed that only the small isoform of the *mocs2* gene, which is the catalytic subunit of the complex, could increase β-cell regeneration. Additionally, sodium molybdate treatment recapitulated the increase in β-cell regeneration observed in our genetic model. Interestingly, sodium molybdate treatment has been shown to lower glucose levels in a mouse model of obesity, in a rat model of β-cell ablation using streptozotocin, as well as in *Drosophila melanogaster* (Perkhulyn et al., 2017; Reul et al., 1997; Tanju Özcelikay et al., 1996). Yet, our results from zebrafish larvae failed to recapitulate the glucose-lowering effect of sodium molybdate treatment, which might be due to dosing or stage. Our data expand on these observations and suggest that perhaps a part of the lower-glucose effect in these models might be due to an increased β-cell mass/functionality.

In both the zebrafish and mouse genetic perturbations of Mocs2, the observed phenotypes were variable between biological replicates suggesting additional factors might be needed to strengthen the β-cell mass and glucose lowering phenotypes. Most of the enzymes that need the molybdopterin paralogues are involved in the regulation of reactive oxygen species (ROS) biology (Sun et al., 2020). ROS have been implicated as regulators of β-cell mass and function in various experimental models. Recently, it was shown that ROS is involved in β-cell proliferation during early postnatal mouse developmental and under conditions that mimic certain aspects of diabetes (Vivoli et al., 2023; Zeng et al., 2017). Similarly, the levels of ROS in the zebrafish larvae were postulated to be important for regulating β-cell proliferation, such that too low or high levels do not stimulate β-cell proliferation (Alfar et al., 2017). Moreover, recent evidence revealed a new molecular mechanism of redox regulation to insulin secretion (Ferdaoussi et al., 2015; Lin et al., 2024; Merrins et al., 2022). Therefore, we hypothesize that the variability in our genetic and chemical models of Moco biosynthesis could stem from a variable level of generated ROS.

Overall, our study describes the liver-to-pancreas axis under conditions of β-cell ablation in the zebrafish. Our genetic screen reveals a previously unexplored role for the molybdenum biosynthetic pathway in β-cell regeneration. Together with previous findings, our observations encourage further exploration of the role of *mocs2* in glucose regulation and diabetes.

## Methods

### Zebrafish transgenic lines

Zebrafish experiments were conducted in compliance with local guidelines and approved by Stockholms djurförsöksetiska nämnd. The previously generated transgenic lines used include: *Tg(ins:flag-NTR)^s950^*(Andersson et al., 2012), *Tg(ins:CFP-NTR)^s892^* (Curado et al., 2007), *Tg(ins:Kaede)^s949^* (Andersson et al., 2012)*, Tg(ins:H2BGFP)^KI112^* (Karampelias et al., 2021), *Tg(fabp10a:Cre)^s955^* (Ni et al., 2012)*, Tg(fabp10a:GFP)^as3^*(Her et al., 2003)*, Tg(fabp10a:CFP-NTR)^s931^* (Choi et al., 2014) *and Tg(-3.5ubb:loxP-EGFP-loxP-mCherry)^cz1701^*(Mosimann et al., 2011) referred to as *ubi:Switch*. As part of this work we generated a stable transgenic line overexpressing the small isoform of *mocs2*, *Tg(fabp10a:smocs2)^KI119^*.

### Mocs2 mouse model generation

The *Mocs2*^+/-^ mouse line (*C57BL/6N-Mocs2^tm1b(EUCOMM)Wtsi^/Ieg*) was constructed using the IMPC ‘knockout first’ targeting strategy at Helmholtz Zentrum München, Germany as follows. Mocs2 mutant mice were generated by allele conversion of C57BL/6N-*Mocs2^tm1a(EUCOMM)Wtsi^/Ieg* mouse line originating from EUCOMM ES clone EPD0560_5_C09 (clone construction overview here: https://www.mousephenotype.org/data/genes/MGI:1336894#order).

Tm1b was produced by crossing *C57BL/6N-Mocs2^tm1a(EUCOMM)Wtsi^/Ieg* mice with an ubiquitously active general Cre deleter mouse line (*C57BL/6NTac-Gt(ROSA)26Sor^tm16(cre)Arte^*) resulting in a deletion of exon 3-5 of Mocs2 (ENSMUSE00001315940; ENSMUSE00001207721; ENSMUSE00001301379) and the neomycin cassette. The mice were genotyped to verify the mutation (genotyping protocol available here: https://infrafrontier.eu/wp-content/uploads/genotype_protocols/EM09099_geno.pdf ).

Heterozygous mice were intercrossed to generate mutant KO mice with wildtype litter mate controls for experimental analysis. There was evidence of postnatal lethality of homozygous offspring. After birth 38% (15 of 50) offspring died. Of remaining offspring 46% *Mocs2*^+/+^ (16 of 35) and 54% *Mocs2*^+/-^ (19 of 35) were present and viable.

The Mocs2 mouse line is available at the European Mouse Mutant Archive (EMMA/Infrafrontier) (https://www.infrafrontier.eu/emma/strain-search/straindetails/?q=9099).

Mice were housed in IVC cages with water and standard mouse chow available ad libitum according to the European Union directive 2010/63/EU and GMC housing conditions (www.mouseclinic.de). All tests were approved by the responsible authority of the district government of Upper Bavaria.

### Mouse experimental procedures

From the age of 8-16 weeks, 14 *Mocs2*^+/-^ mice (7m/7f) and *Mocs2*^+/+^ controls (7m/7f) were phenotyped systematically in the German Mouse Clinic (GMC) as described previously (Fuchs et al., 2011; Fuchs et al., 2018) and in accordance with the standardised phenotyping pipeline of the IMPC (IMPReSS: https://www.mousephenotype.org/impress/index).

Altered glucose metabolism was examined using the intraperitoneal glucose tolerance test (ipGTT) at the age of 13 weeks. Glucose was administered intraperitoneally (2 g/kg i.p.) after a 16-h withdrawal of food with body weight being determined before and after food withdrawal and glucose levels being measured before and at 15, 30, 60, and 120 minutes after glucose injection. Blood glucose concentrations were assessed in blood collected from the tail vein with the Accu-Chek Aviva Connect glucose analyzer (Roche/Mannheim).

At the age of 16 weeks, the final blood samples were collected from the retrobulbar vein plexus under isoflurane anesthesia in Li-heparin coated tubes (Li1000A, Kabe Labortechnik). The samples were centrifuged at 5000xg for 10 minutes at 8°C and plasma separated within two hours after blood collection. Clinical chemistry parameters were measured immediately using an AU480 analyzer (Beckman-Coulter) and adapted reagent kits from Beckman-Coulter according to the manufacturer’s instructions, as described previously (Rathkolb et al., 2013).

Body weight was taken weekly prior to the different testing procedures throughout the whole phenotyping pipeline procedure. Tissues derived from wild-type and mutant mice were fixed in neutral buffered formalin, processed, embedded in paraffin, sectioned at 3 µm and stained with a standard hematoxylin and eosin protocol. The double immunohistochemical staining was performed using a BOND RX^m^ (Leica, Germany) automated stainer. Pancreas sections were deparaffinized, antigens were retrieved with citrate buffer for 30 minutes and blocked for 30 minutes with blocking agent. Primary antibodies (insulin rabbit monoclonal, Cell Signaling 3014, 1:3000 and glucagon, mouse monoclonal, Sigma-Aldrich G2654, 1:1000) were applied and incubated for 60 minutes followed by secondary antibodies. The detection was performed with DAB and Red BOND polymers. Slides were counterstained with hematoxylin and mounted with a DAKO mounting medium. The slides were imaged with a NanoZoomer S60 (Hamamatsu, Japan) scanner. For the immunofluorescence procedure, sections were deparaffinized, rehydrated and antigens were retrieved with Sodium Citrate pH 6 buffer by microwave heating for 15 minutes. Sections were permeabilized with 0.3% triton-X100, followed by blocking solution and primary antibody incubation with anti-urocortin3 (rabbit, Phoenix Pharmaceuticals H-019-29, 1:300) and anti-glucagon (mouse, Santa Cruz sc-514592, 1:400) overnight, followed by a 2-hour incubation at room temperature with fluorescent conjugated secondary antibodies. Imaging was performed using a Leica SP5 confocal microscope.

### Chemical treatments of zebrafish larvae

β-cell and hepatocyte ablation in the *Tg(ins:CFP-NTR)*, *Tg(ins:flag-NTR)* or *Tg(fabp10a:CFP-NTR)* was performed by incubating the zebrafish for 24 hours with 1 mM (2-month-old fish) or 10 mM (larvae) MTZ (Sigma-Aldrich) diluted in 1% DMSO (VWR) in facility water (2-month-old fish) or an E3 solution supplemented with 0.2 mM 1-phenyl-2-thiourea (PTU; Acros Organics) (larvae). Hepatocyte injury was performed by incubating the zebrafish larvae with 10 mM acetaminophen (Sigma-Aldrich) for 24 hours. Sodium molybdate (Sigma-Aldrich) was added to the zebrafish water at a final concentration of 10 μM.

### Immunostaining of zebrafish larvae

Immunofluorescence of zebrafish larvae was described previously (Lu et al., 2016). Primary antibodies were used against GFP (1:500; Aves Labs GFP-1020), insulin (1:100; custom made by Cambridge Research Biochemicals) and somatostatin 1:300 (DAKO-A0566). Liver and insulin area was calculated on maximum projections using the Fiji threshold function.

### Glucose measurements in zebrafish larvae

Glucose levels were measured using the Glucose Colorimetric/Fluorometric Assay kit (BioVision) in pools of 4 larvae for each time point and condition. Blood glucose measurements in 2-month-old zebrafish were performed using a standard glucometer (Freestyle – Abbott). Fish were fasted for 16 h, anesthetized in tricaine (Sigma-Aldrich) and decapitated for blood glucose measurements.

### Hepatocyte isolation and RNASeq

Three independent hepatocyte isolations from 2-month-old zebrafish were performed from three different crosses. β-cells were ablated with MTZ for 24 hours followed by a washout of the drug, feeding and regeneration for an additional day. Zebrafish treated with MTZ but without the *Tg(ins:flag-NTR)* transgene were used as controls to adjust for the effect of the MTZ on the transcriptome. 10-15 zebrafish of mixed sex were sacrificed by head decapitation and the livers were dissected with the help of a fluorescent microscope, as the fish carried the *Tg(fabp10a:GFP)* transgene. Then livers were cut into smaller pieces, passed through a needle to further disrupt the tissue and incubated with TrypLE for 45 min at 37 degrees Celsius until a single suspension was created. TrypLE was inactivated using fetal bovine serum and the single suspension was sorted using the FACSAria III instrument and the FITC-A filter to sort the GFP^+^ cells.

RNA was extracted from the sorted cells using the RNAqueous micro total isolation kit (Invitrogen). The quality of the RNA was assessed and the first control sample was discarded as the RIN value was too low. Libraries were prepared using the TRUSeq stranded mRNA kit (Illumina) and sequenced on an Illumina HiSeq2500 instrument. Reads were mapped using Tophat 2.0.4 to the GRCz10 zebrafish genome assembly. Differential expression analysis was conducted using the DESEQ package on the R environment (Anders and Huber, 2010). Gene ontology analysis was performed using the Panther online database (version 15.0) (Mi et al., 2009; Thomas et al., 2003).

### Cloning of selected genes and genetic screen

Cloning of *sdf2l1* and the selected enzymes was performed using the Gateway system. Of note, we did not manage to clone two significantly upregulated enzymes, namely *pdzrn3a* and *tor1l1* due to a lack of cDNA amplification. Pooled cDNA from different developmental zebrafish larval stages were used as template to amplify the gene sequences, which were subsequently cloned into the pDONR221 middle entry vector. Primer sequences for the amplification of the genes are shown in Supplementary Table 1. We also generated a 5’ entry vector containing the *fabp10a* promoter. Then we performed three-way recombination reactions using the 5’ entry clone containing the *fabp10a* promoter, the middle entry vectors containing the different genes, the 3’ *polyA* entry vector and the pDESTtol2CG2 destination vector that carries the *cmcl2:GFP* transgene for selection. The final vectors were sequenced and then 15 pg of the vectors together with 20 pg of transposase mRNA were injected into the one-cell-stage zebrafish embryos to induce mosaic overexpression. At 3 dpf the larvae were screened for the transgene integration by assessing the presence of the *cmcl2:GFP* expression. The regeneration assay was performed by ablating the β-cells between 3-4 dpf and manually counting the number of β-cells using a fluorescence microscope at 6 dpf.

### Expression analysis using publicly available scRNA-Seq datasets

For the mouse pancreas gene expression analysis: we used the processed and annotated mouse pancreas scRNA-Seq atlas dataset (Hrovatin et al., 2023) which was downloaded from GEO with accession number: GSE211799, file name: GSE211799_adata_atlas.h5ad.gz. The .h5ad file was imported into a jupyter notebook and violin plots were generated using the Scanpy toolkit (1.9.5) and the ”cell_type_integrated_v2_parsed” clustering option (Wolf et al., 2018).

For the human liver gene expression analysis: we used the processed and annotated human liver pancreas scRNA-Seq atlas .h5ad file, downloaded from the pre-print associated website (https://liver.unifiedcellatlas.org), file name: ’normal_final_webV2.h5ad’ (Wu et al., 2023). We visualized the expression using violin plots with the ’level 1’ obs clustering option of the file in the Scanpy toolkit (1.9.5).

For the human islet dataset: we used the processed and annotated human islet scRNA-Seq dataset from (Elgamal et al., 2023) which was downloaded from (https://www.gaultonlab.org/pages/Islet_expression_HPAP.html). The file was imported in Rstudio using R version 4.2.3 (Team, 2019) and the violin plots were generated with the group.by=”Cell Type” annotation using the Seurat package 4.4.0 (Hao et al., 2021).

### Statistical analysis

Statistical analysis throughout the manuscript was carried out using the GraphPad Prism software. *P* values ≤ 0.05 were considered significant. Normal distribution of data was calculated using a combination of the Shapiro-Wilk and D’Agostino&Pearson tests. *P* values are reported in the respective figure legends of the experiments where appropriate.

---

**Supplementary Table 1:**
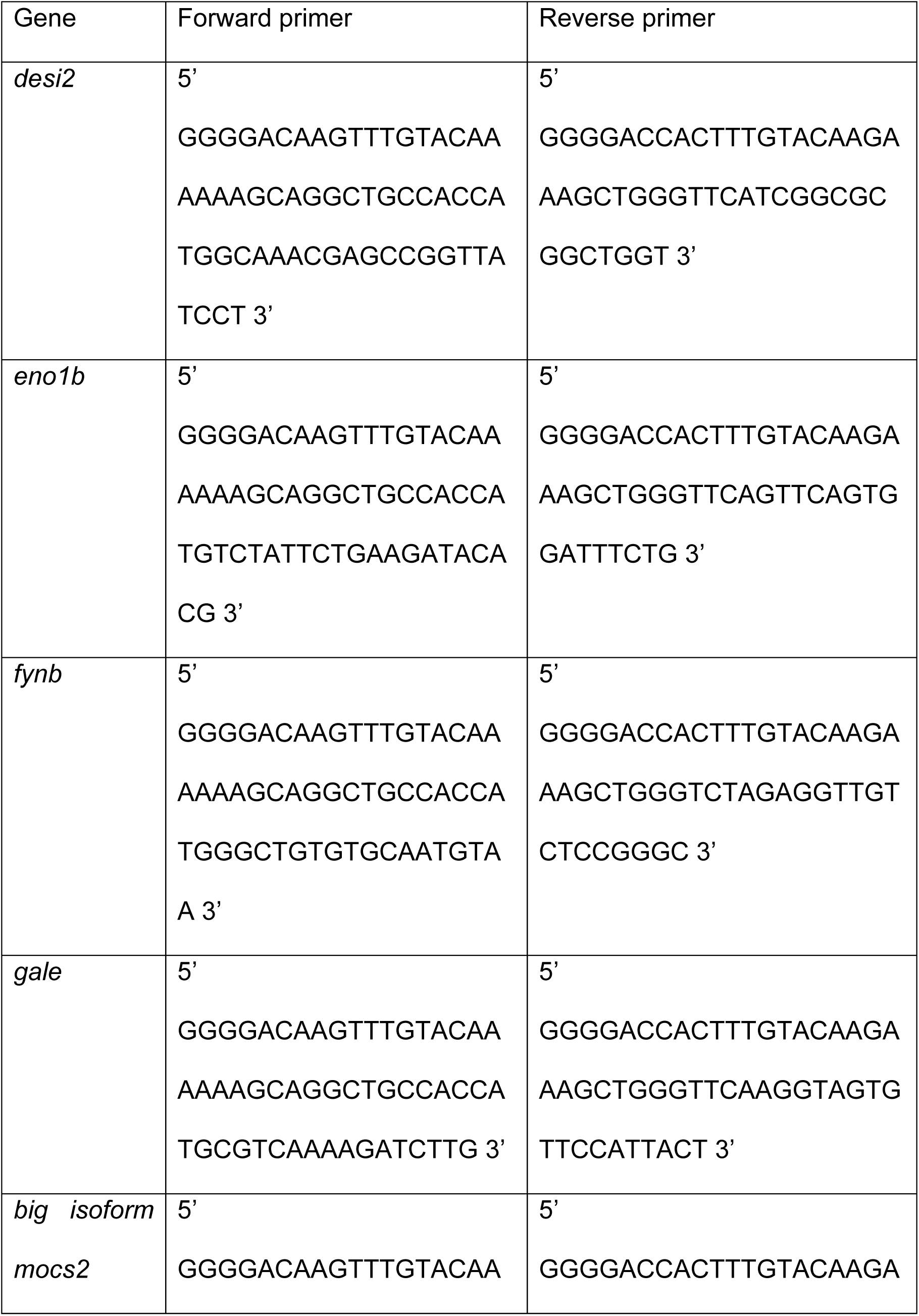
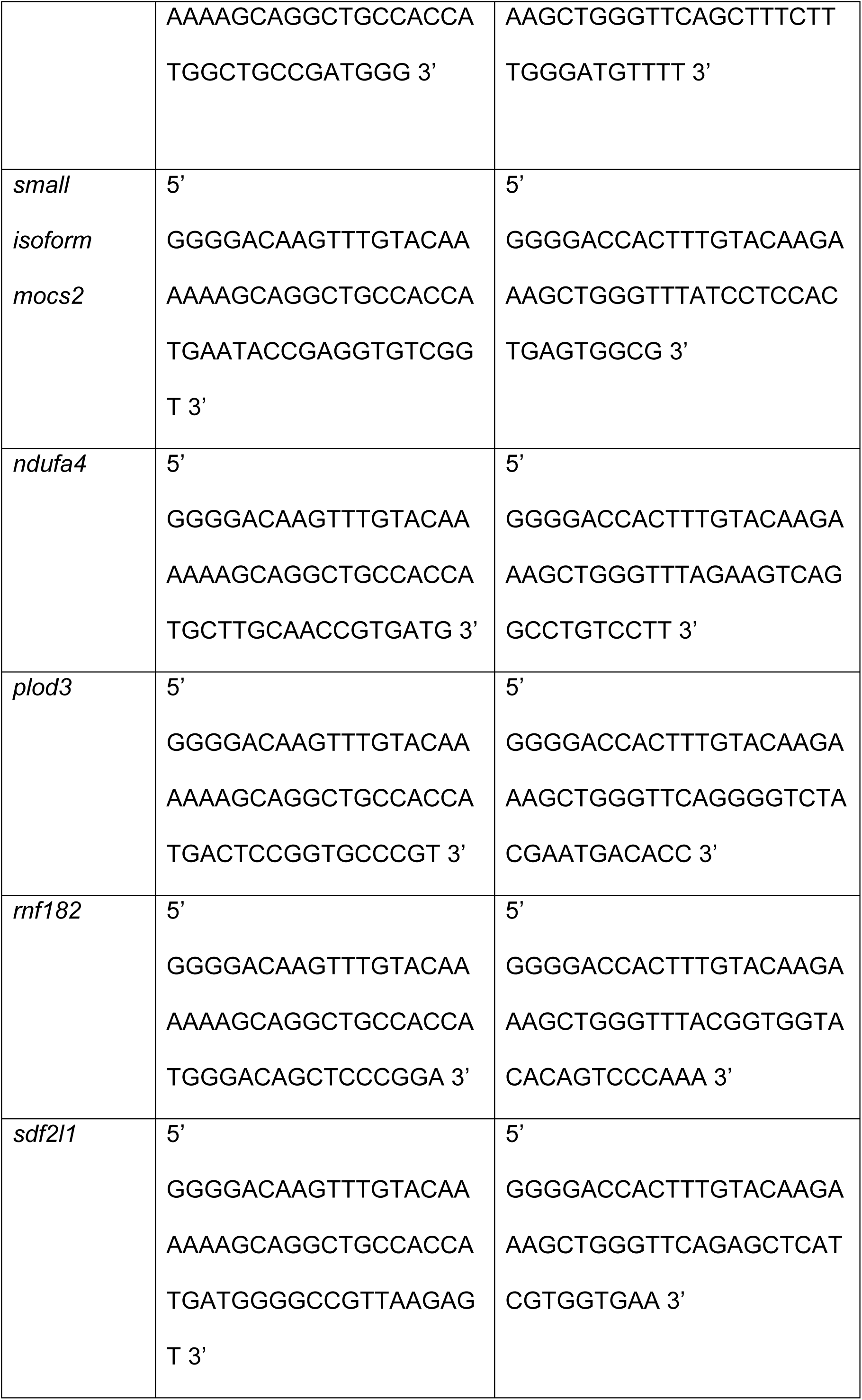
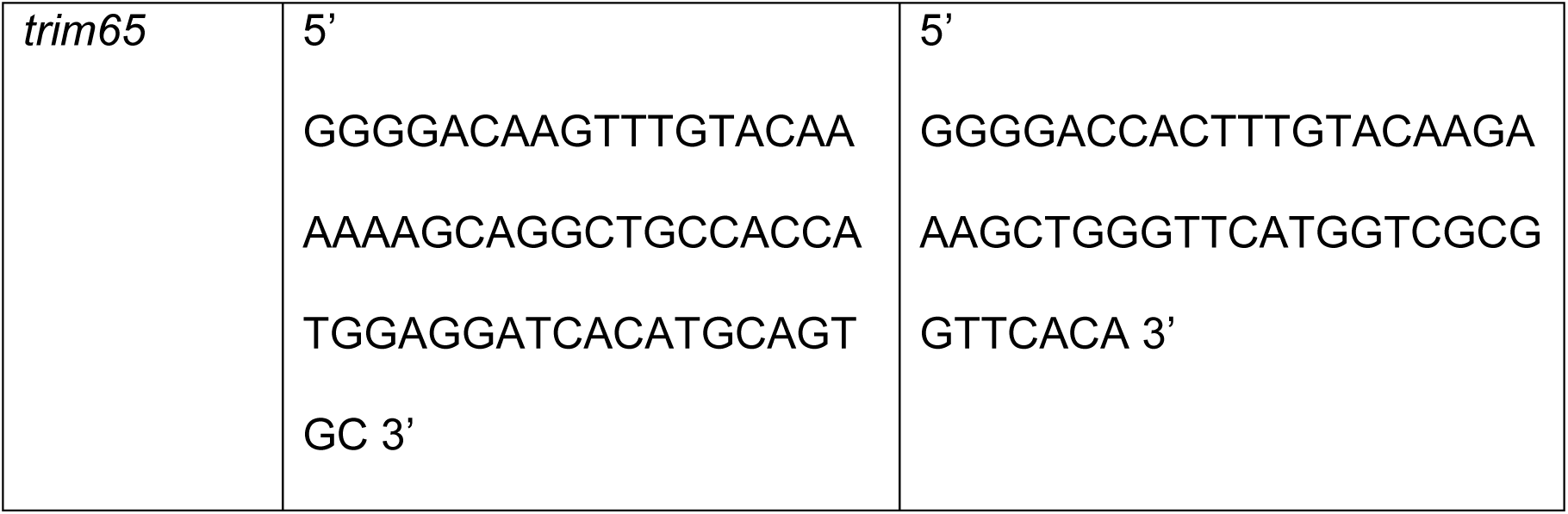
Primers used for amplification of selected genes to clone into the entry vectors of the Gateway system.

## Data availability

The data that support the findings of this study are available from the corresponding authors upon reasonable request. Raw RNA-Seq data will become available upon publication.

## Acknowledgments

Research in the lab of O.A. was supported by funding from the Swedish Research Council, the Novo Nordisk Foundation, Diabetes Wellness, Cancerfonden, and the Strategic Research Programmes in Diabetes at the Karolinska Institutet. E.B.W. was funded by a postdoctoral fellowship from Novo Nordisk A/S. The German Mouse Clinic was supported by German Federal Ministry of Education and Research (Infrafrontier grant 01KX1012 to MHdA); German Center for Diabetes Research (DZD) (MHdA).

## Competing interests

The authors declare no competing interests.

## Author contributions

C.K designed, performed and analyzed most of the experiments in the study. E.B.W performed experiments with the *sdf2l1* construct. B.B and L.C performed zebrafish experiments. B.R and P.d.S-M designed, performed and analyzed mouse data. S.M generated the mouse model. H.F and V.G-D managed and supervised the mouse project. M.H.d.A acquired funding for and supervised the mouse project. O.A designed, supervised and acquired funding for the study. C.K and O.A wrote the manuscript with input from all authors.

## Supplementary Figure Legends

**Supplementary Figure 1:**
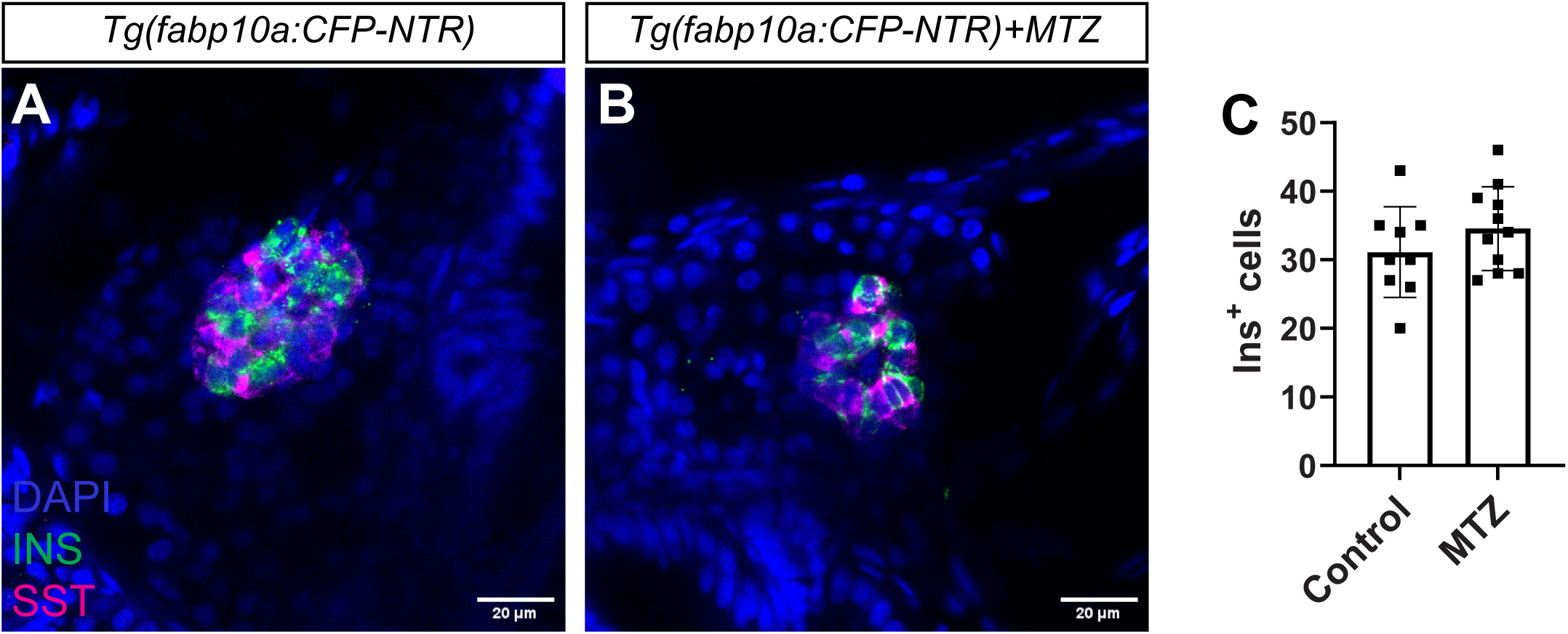
Hepatocyte ablation does not affect β-cell development. A-C: Single-plane confocal images of pancreatic islets from *Tg(fabp10a:CFP-NTR)* larvae treated with DMSO (**A**) or MTZ (**B**) for 24 hours. Quantification of β-cells (INS^+^ in green) (**C**) did not show any difference between the conditions. Scale bar 20 µm. *n*=9-11.

**Supplementary Figure 2:**
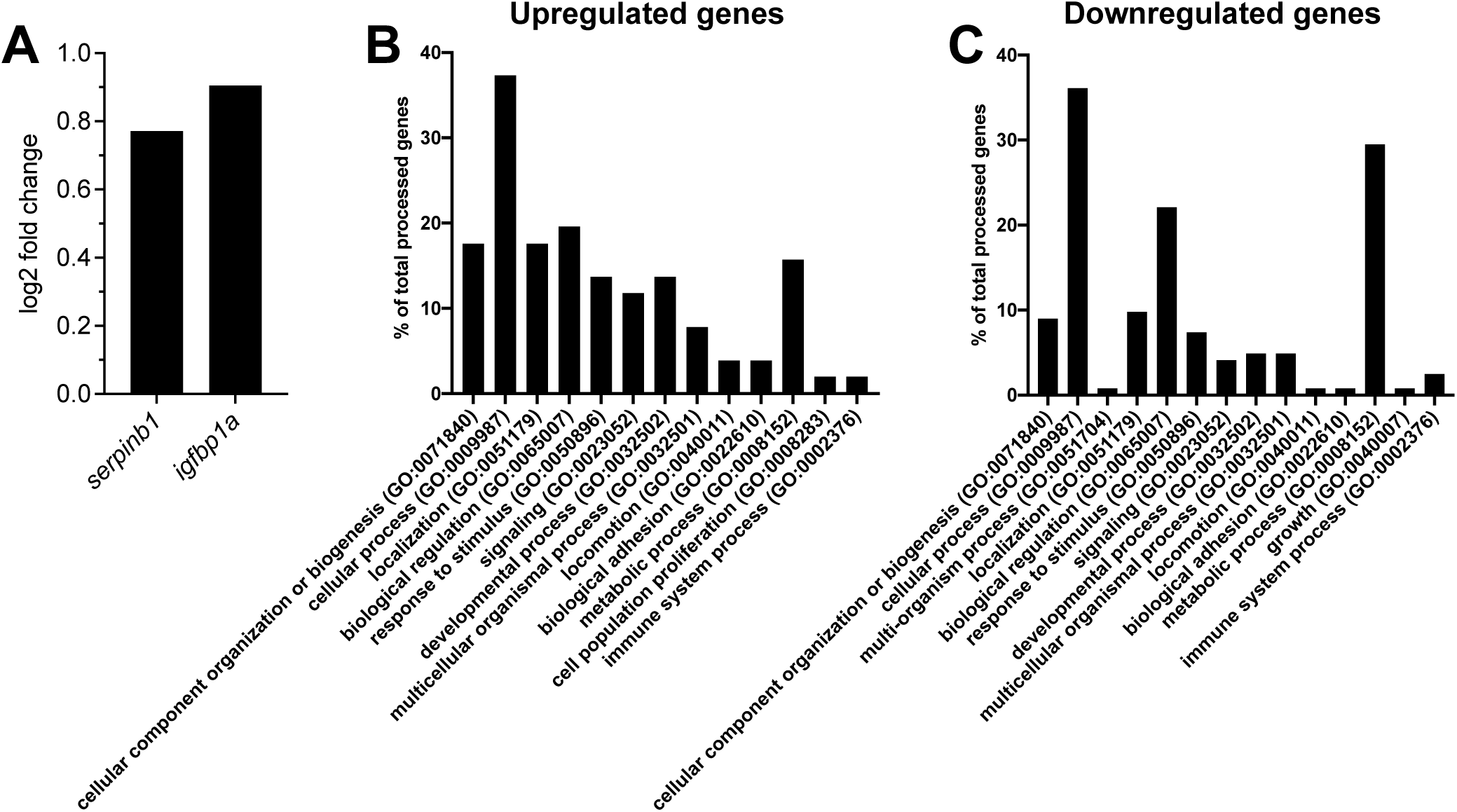
Pathway enrichment analysis of RNASeq dataset. **A**: Log2 fold change levels of *serpinb1* and *igfbp1a* in hepatocytes following β-cell ablation. **B**-**C**: Enriched pathways for the upregulated (**B**) and downregulated (**C**) genes following β-cell ablation in the RNASeq dataset.

**Supplementary Figure 3:**
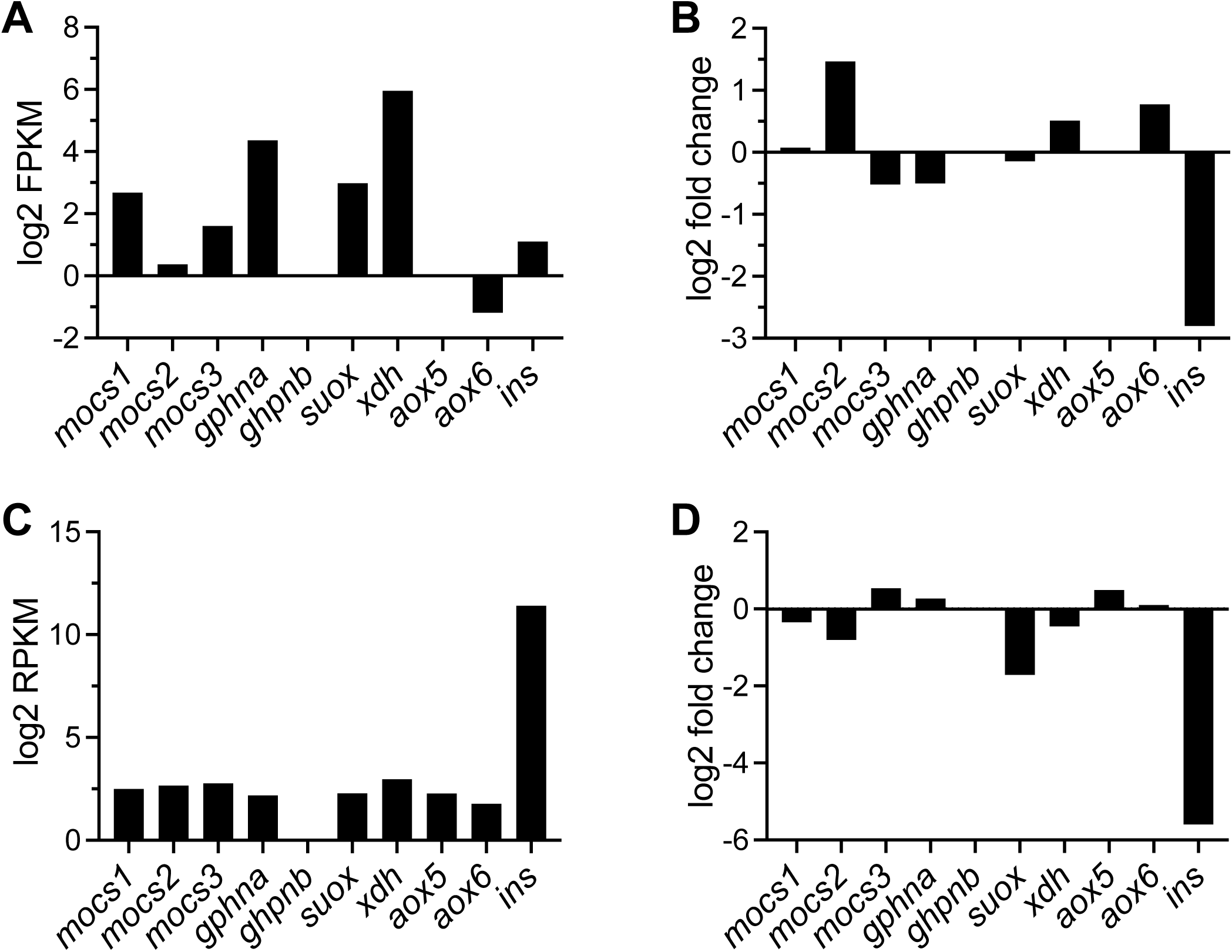
Expression levels of genes of the Moco biosynthetic pathway in hepatocytes and pancreatic islets from zebrafish. **A**-**B**: Expression levels of the enzymes involved in Moco biosynthesis and utilization in isolated hepatocytes (**A**). Fold change of the same enzymes in hepatocytes following β-cell ablation (**B**). **C**-**D**: Expression levels of the enzymes involved in Moco biosynthesis and utilization in isolated islets from zebrafish larvae (**C**). Fold change of the same enzymes in primary islets of zebrafish larvae following β-cell ablation (**D**).

**Supplementary Figure 4:**
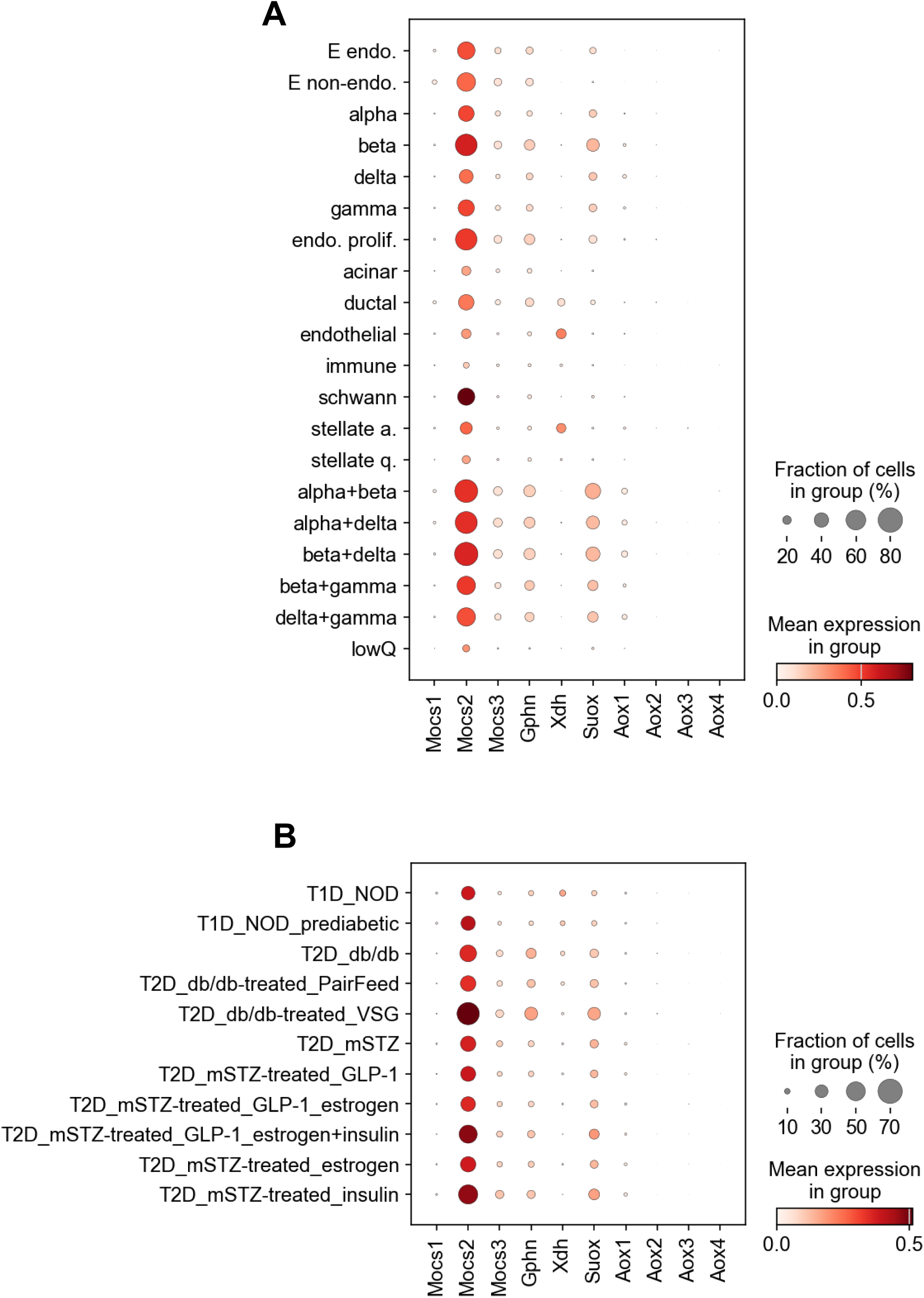
Expression levels of genes involved in the Moco biosynthetic pathway in mouse pancreas. **A-B**: Dot plots showing the expression levels of the core gene of the Moco biosynthetic pathway across the different cell populations of the mouse pancreas (**A**) and across different models of diabetes in mice (**B**).

**Supplementary Figure 5:**
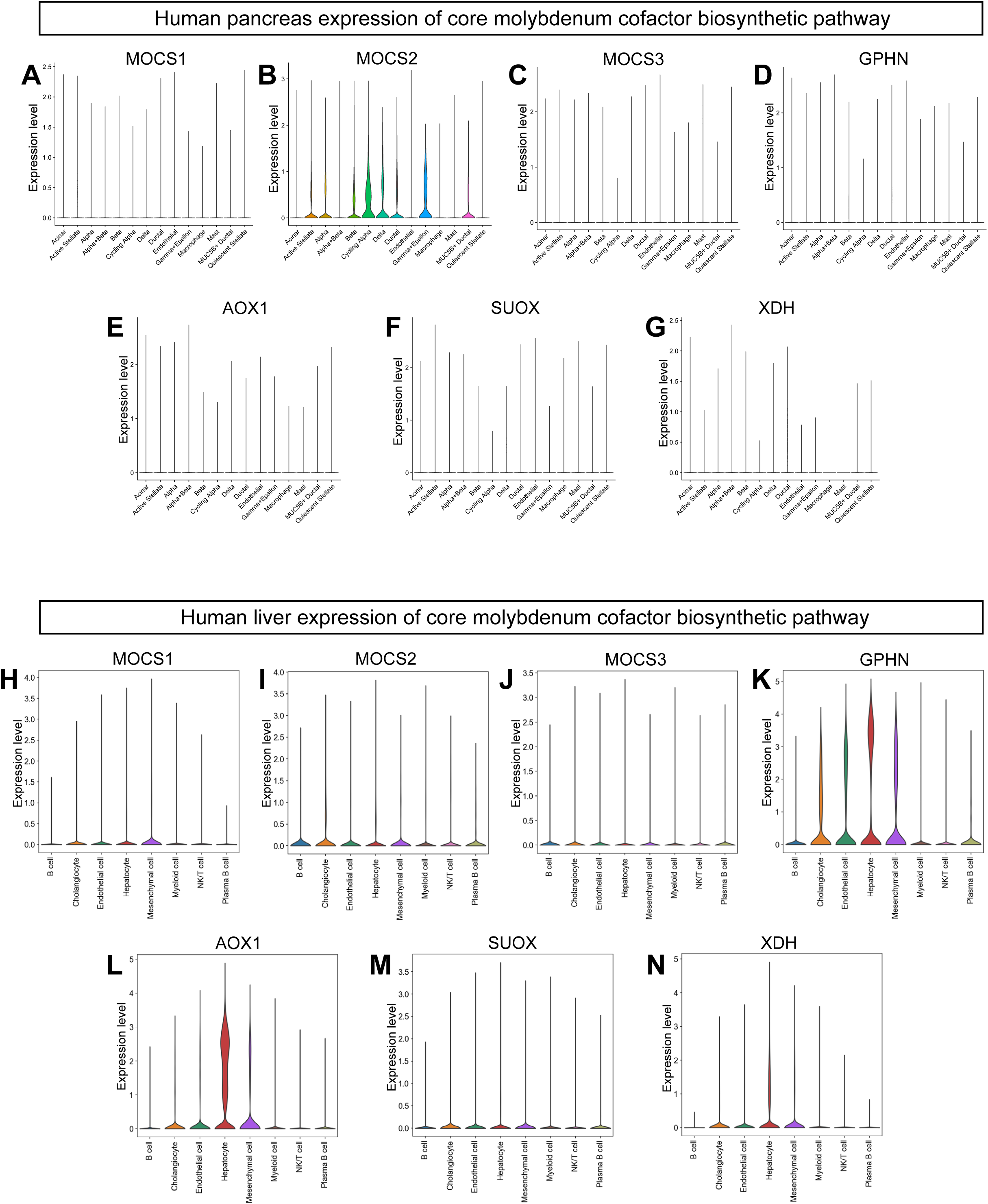
Expression levels of genes involved in the Moco biosynthetic pathway in human pancreas and liver. **A-G**: Violin plots showing the expression levels of the core gene of the Moco biosynthetic pathway across the different cell populations of the human pancreas. **H-N**: Violin plots showing the expression levels of the core gene of the Moco biosynthetic pathway across the different cell populations of the human liver.

**Supplementary Figure 6:**
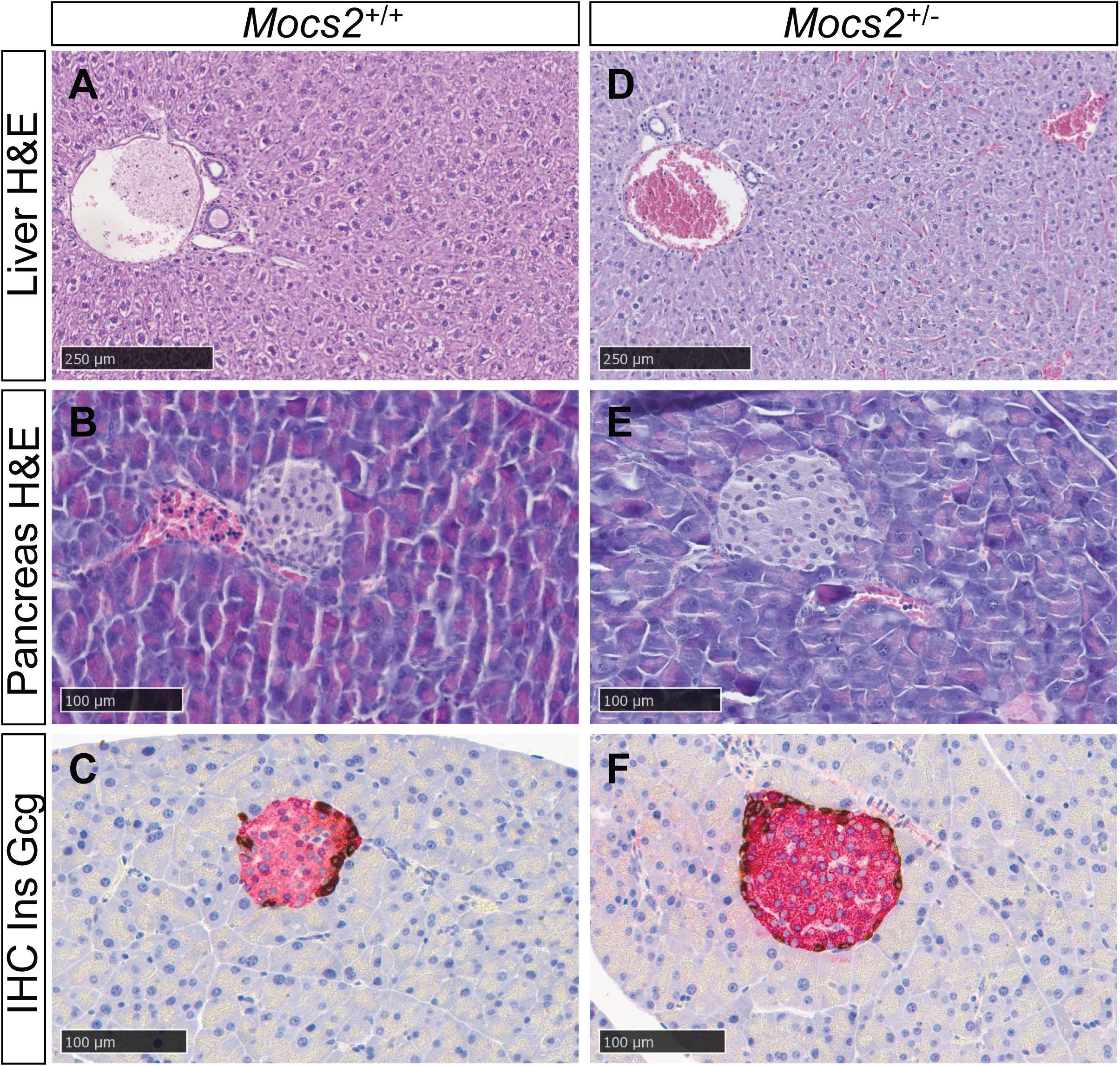
Hematoxylin and eosin photomicrographs of liver and pancreas and double immunohistochemistry images of pancreas of *Mocs2*^+/-^ mice. **A-F**: Images showing no abnormalities or differences in any examined tissue between *Mocs2*^+/+^ and *Mocs2*^+/-^ male mice. Representative H&E brightfield images from liver and pancreas from *Mocs2*^+/+^ **(A-B)** und *Mocs2*^+/-^ **(D-E)** male mice. Immunohistochemical double staining of endocrine pancreas to assess islet morphology, showing insulin-(Ins, red) and glucagon-producing cells (Gcg, brown) in *Mocs2*^+/+^ (**C**) und *Mocs2*^+/-^ (**F**) male mice. The morphology was similar to that of female mice. Scale bars are indicated in the figure.

**Supplementary Figure 7:**
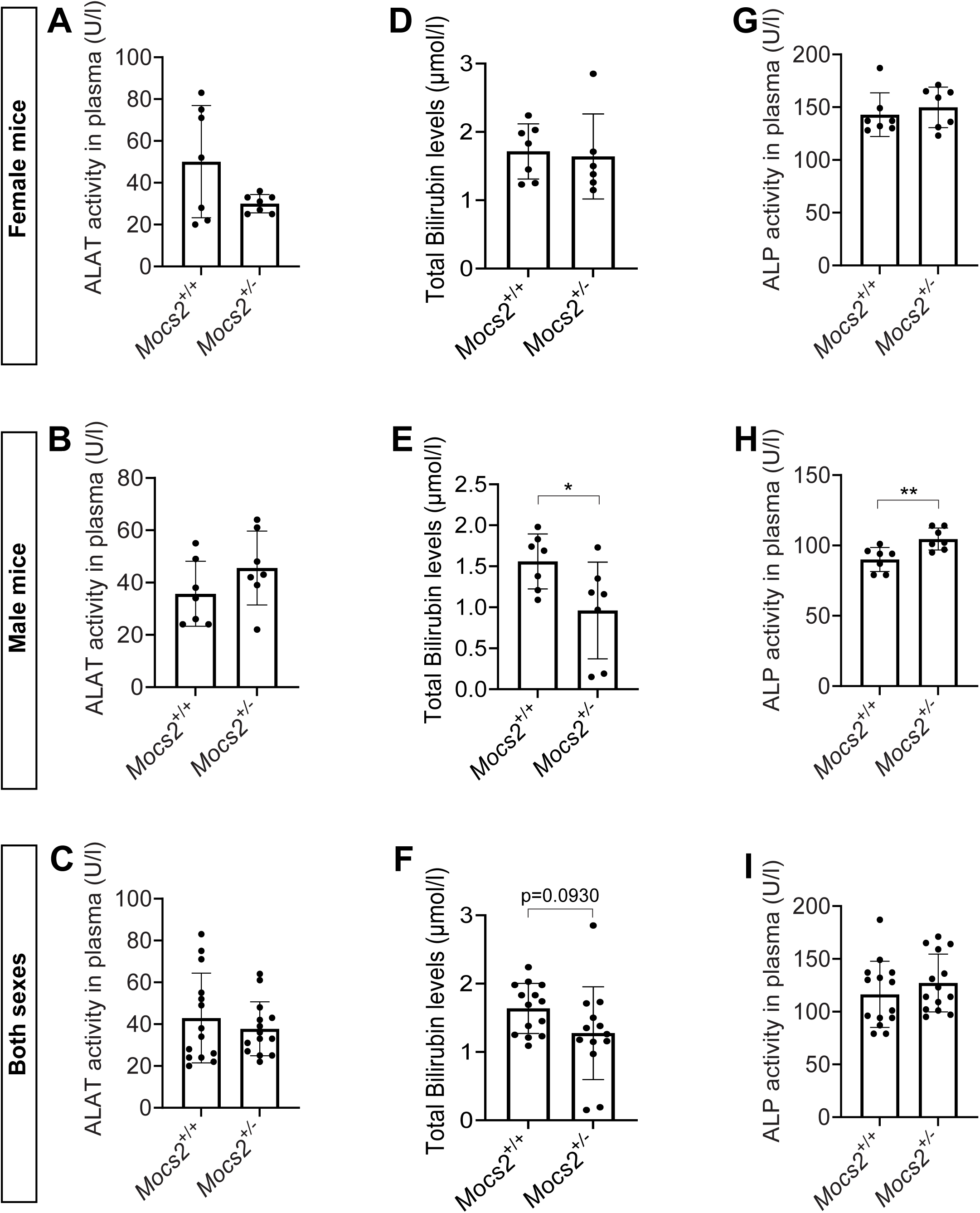
Clinical markers of liver functionality of *Mocs2^+/-^* mice. A-C: ALAT activity (**A**), Bilirubin levels (**B**) and ALP activity (**C**) in plasma of female *Mocs2^+/+^* and *Mocs2^+/-^*mice. **D-F:** ALAT activity (**D**), Bilirubin levels (**E**) and ALP activity (**F**) in plasma of male *Mocs2^+/+^* and *Mocs2^+/-^*mice. **G-I:** ALAT activity (**G**), Bilirubin levels (**H**) and ALP activity (**I**) in plasma of combined male and female *Mocs2^+/+^*and *Mocs2^+/-^*mice.

## Notes

### Competing Interest Statement

The authors have declared no competing interest.

